# Accurate Predictions of Postmortem Interval Using Linear Regression Analyses of Gene Meter Expression Data

**DOI:** 10.1101/058370

**Authors:** Colby M. Hunter, Alex E. Pozhitkov, Peter A. Noble

## Abstract

In criminal and civil investigations, postmortem interval is used as evidence to help sort out circumstances at the time of human death. Many biological, chemical, and physical indicators can be used to determine the postmortem interval, but most are not accurate. Here, we sought to validate an experimental design to accurately predict the time of death by analyzing the expression of hundreds of upregulated genes in two model organisms, the zebrafish and mouse. In a previous study, the death of healthy adults was conducted under strictly controlled conditions to minimize the effects of confounding factors such as lifestyle and temperature. A total of 74,179 microarray probes were calibrated using the Gene Meter approach and the transcriptional profiles of 1,063 significantly upregulated genes were assembled into a time series spanning from life to 48 or 96 h postmortem. In this study, the experimental design involved splitting the gene profiles into training and testing datasets, randomly selecting groups of profiles, determining the modeling parameters of the genes to postmortem time using over- and/or perfectly- defined linear regression analyses, and calculating the fit (R^2^) and slope of predicted versus actual postmortem times. This design was repeated several thousand to million times to find the top predictive groups of gene transcription profiles. A group of eleven zebrafish genes yielded R^2^ of 1 and a slope of 0.99, while a group of seven mouse liver genes yielded a R^2^ of 0.98 and a slope of 0.97, and seven mouse brain genes yielded a R^2^ of 0.93 and a slope of 0.85. In all cases, groups of gene transcripts yielded better postmortem time predictions than individual gene transcripts. The significance of this study is two-fold: selected groups of upregulated genes provide accurate prediction of postmortem time, and the successfully validated experimental design can now be used to accurately predict postmortem time in cadavers.

## Introduction

The postmortem interval (PMI) is the elapsed time between death of an organism and the initiation of an official investigation to determine the cause of death. Its determination is important to civil investigations such as those involving life insurance fraud because investigators need to determine if the person was alive or not when the policy was in effect [1]. The PMI is also important to criminal investigations, especially suspicious death cases where there are no witnesses, because it can help determine the time relationship between a potential suspect and the victim and eliminate people from a suspect list, which speeds up investigations. Accurate prediction of PMI is considered one of the most important and complex tasks performed by forensic investigators [2].

Several studies have suggested that RNA could be used to estimate PMI [3,4,5,6,7]. While most studies focused on the degradation of mRNA gene markers, some examined gene expression. The RNA degradation studies include: a model to predict PMI based on the degradation of Beta actin (*Actb*), Glyceraldehyde-3-phosphate dehydrogenase (*Gapdh*), Cyclophilin A (*Ppia*) and Signal recognition particle 72 (*Srp72*) genes in mouse muscle tissue samples [3], a model to predict PMI based on degradation of an amplified *Actb* gene and temperature in rat brain samples [4], and a study that predicted PMI based on the degradation of *Gapdh, Actb* and 18S rRNA genes in the spleens of rats [5]. The gene expression studies include: a study that found increased expression of myosin light chain 3 (*Myl3*), matrix metalloprotease 9 (*Mmp9*) and vascular endothelial growth factor A ( *Vegfa*) genes in human body fluids after 12 h postmortem [6], a study that found increased expression of Fas Ligand (*Fasl*) and ‘phosphatase and tensin homologue deleted on chromosome 10’ (*Pten*) genes with postmortem time in rats [7], and a study that found individual gene transcripts did not increase using PCR-based gene expression arrays of frozen human brain cadaver samples [8]. Common to these studies is the requirement: (i) to amplify cDNA by polymerase chain reaction (PCR) and (ii) to normalize the data with a control in order to facilitate sample comparisons. These requirements introduce methodological biases that could significantly affect interpretation of the data. An approach that minimizes or eliminates these biases is highly desirable because it might lead to better PMI predictions.

Since conventional DNA microarray approaches yield noisy data [9], in 2011 we developed the “Gene Meter” approach that precisely determines specific gene abundances in biological samples and minimizes noise in the microarray signal [10,11]. The reason this approach is precise is because the behavior of every microarray probe is determined by calibration - which is analogous to calibrating a pH meter with buffers. Without calibration, the precision and accuracy of a meter is not known, nor can one know how well the experimental data fits to the calibration (i.e., R). The advantages of the Gene Meter approach over conventional DNA microarray approaches is that the calibration takes into consideration the non-linearity of the microarray signal and calibrated probes do not require normalization to compare biological samples. Moreover, PCR amplification is not required. We recognize that next-generation-sequencing (NGS) approaches could have been used to monitor gene expression in this study. However, the same problems of normalization and reproducibility are pertinent to NGS technology [12]. Hence, the Gene Meter approach is currently the most advantageous high throughput methodology to study postmortem gene expression and might have utility for determining the PMI.

The Gene Meter approach has been used to examine thousands of postmortem gene transcription profiles from 44 zebrafish (*Danio reno*) and 20 house mice (*Mus musculus*) [13]. Many genes were found to be significantly upregulated (relative to live controls). Given that each sampling time was replicated two or three times, we conjectured that the datasets could be used to assess the feasibility for predicting PMIs from gene expression data. Although many approaches are available to determine PMI (see Discussion), an approach that accurately determines the time of death is highly desired and it is the goal of our study to determine if specific gene transcripts or groups of gene transcripts could accurately predict postmortem time. Zebrafish and mice are ideal for testing experimental designs because the precise time of human deaths is often not known, and other variables such as lifestyle, temperature, and health condition are also often not known or sufficiently controlled in human studies. Given that these variables could have confounding effects on the interpretation of gene expression data in human studies, testing experimental designs under controlled conditions using model organisms is ideal. In our study, the timing of death and health of the zebrafish and mice are known, which enables the testing of different experimental designs to provide “proof of principle”. It is our intent to use the best design to determine PMI of cadavers for future studies.

The objectives of our study are twofold: (1) to identify specific upregulated genes or groups of upregulated genes that accurately predict the PMI in the zebrafish and mouse, and (2) to design and evaluate a robust experimental approach that could later be implemented to predict PMI from cadavers.

## Materials and Methods

Although the details of zebrafish and mouse processing, the extraction of RNA, and microarray calibrations are presented in a previous study [13], we have provided relevant experimental protocols to aid readers in the interpretation of the results of this study.

**Zebrafish processing**. The 44 zebrafish were maintained under standard conditions in flow-through aquaria with a water temperature of 28°C. Prior to sacrifice, the zebrafish were placed into 1 L of water of the same temperature as the aquaria. At zero time, four fish were extracted and snap frozen in liquid nitrogen. These live controls were then placed at −80°C. To synchronize the time of death, the remaining 40 fish were put into a small container with a bottom made of mesh and placed into an 8 L container of ice water for 5 mins. The small container with the mesh bottom was placed into the flow-through aquarium with a water temperature of 28°C for the duration of each individual’s designated postmortem time. The postmortem sampling times used for the zebrafish were: 0, 15 min, 30 min, 1, 4, 8, 12, 24, 48 and 96 h. At each sampling time, 4 individuals were taken out of the small container in the flow-through aquarium, snap frozen in liquid nitrogen and then stored at −80°C. One zebrafish sample was not available for use (it was accidentally flushed down the sink) however this was taken into account for calculation of extraction volumes.

**Mouse processing**. Twenty C57BL/6JRj male mice of the same age (5 months) and similar weight were used. Prior to sacrifice, the mice had *ad libitum* access to food and water and were maintained at room temperature. At zero time, the mice were sacrificed by cervical dislocation and each mouse was placed in a unique plastic bag with pores to permit the transfer of gases. The mice were kept at room temperature for the designated postmortem sampling times. The sampling times used were: “zero” time, 30 min, 1, 6, 12, 24 and 48 h. At each sampling time, a brain and two liver samples were obtained from each of three mice, except for the 48 h sampling where only two mice were sampled. The samples were immediately snap frozen in liquid nitrogen and placed at −80°C.

**RNA Processing and Labeling**. Gene expression samples for each PMI were done in duplicate for zebrafish and in triplicate for mice (except for the 48 h PMI sample that was duplicated). The zebrafish samples were homogenized with a TissueLyzer (Qiagen) with 20 ml of Trizol. The mouse brain and liver samples (~100 mg) were homogenized in 1 ml of Trizol. One ml of the homogenate was placed into a centrifuge tube containing 200 |il of chloroform. The tube was vortexed and placed at 25°C for three min. Following centrifugation for 15 min at 12000 RPM, the supernatant was placed into a new centrifuge tube containing an equal volume of 70% ethanol. Purification of the RNA was accomplished using the PureLink RNA Mini Kit (Life Technologies, USA). The purified RNA was labeled using the One-Color Microarray-based Gene Expression Analysis (Quick Amp Labeling). The labeled RNA was hybridized to the DNA microarrays using the Tecan HS Pro Hybridization kit (Agilent Technologies). The zebrafish RNA was hybridized to the Zebrafish (v2) Gene Expression Microarray (Design ID 019161) and the mouse RNA was hybridized to the SurePrint G3 Mouse GE 8x60K Microarray Design ID 028005 (Agilent Technologies) following the manufacturer’s recommended protocols. The microarrays were loaded with 1.65 μg of labeled cRNA for each postmortem time and sample.

**Calibration of the DNA microarray**. Oligonucleotide probes were calibrated by hybridizing pooled serial dilutions of all samples for the zebrafish and the mouse. The dilution series for the Zebrafish array was created using the following concentrations of labeled cRNA: 0.41, 0.83, 1.66, 1.66, 1.66, 3.29, 6.60, and 8.26 μg. The dilution series for the Mouse arrays was created using the following concentrations of labeled cRNA: 0. 17, 0.33, 0.66, 1.32, 2.64, 5.28, 7.92, and 10.40 μg. The behavior of each probe was determined from these pooled dilutions as described in the previous studies [10,11]. The equations of the calibrated probes were assembled into a dataset so that they could be used to back-calculate gene abundances of unknown samples (Supporting Information Files S1 and S2 in Ref. 13).

**Statistical analyses**. Gene transcription profiles were constructed from the gene abundance data determined from the 74,179 calibrated profiles. Expression levels were log-transformed for analysis to stabilize the variance. A one-sided Dunnett’s T-statistic was applied to test for upregulation at one or more postmortem times compared to live control (fish) or time 0 (mouse). A bootstrap procedure with 10^9^ simulations was used to determine the critical value for the Dunnett statistics in order to accommodate departures from parametric assumptions and to account for multiplicity of testing. The profiles for each gene were centered by subtracting the mean values at each postmortem time point to create “null” profiles. Bootstrap samples of the null profiles were generated to determine the 95th percentile of the maximum (over all genes) of the Dunnett statistics. Significant postmortem upregulated genes were selected as those having Dunnett T values larger than the 95th percentile. Only significantly upregulated genes were retained for further analyses. The significantly upregulated transcriptional profiles are found in the Supporting Information - Compressed/ZIP File Archive. The archive contains 3 files: zebrafish_calib_probe_abundance.txt, mice_liver_probe_log10_abundance.txt, and mice_brain_probe_log10_abundance.txt. Each file has the following four columns: Agilent Probe Identification Tag, sample time, sample number and log10 concentration.

The software for calculating the numerical solution of the over- and perfectly-defined linear regressions was coded in C++ and has been used in previous studies [14,15]. The C++ code allowed us to train and test thousands to millions of regression models. A description of the analytical approach can be found in the original publication [15]. Briefly, the abundances of each gene transcript in a gene set was numerically solved in terms of predicting the postmortem times with modeling parameters (i.e. coefficients). A version of the C++ source code is available at http://peteranoble/software under the heading: “Determine the coefficients of anxs equation using matrix algebra”. The web page includes a Readme and example files to help users implement the code. To aid readers in understanding the linear matrix algebra used in the study, we have provided a primer in the Supplemental Information section. The postmortem time was predicted from the sum of the product of the gene abundances multiplied by the coefficients for each gene transcript. Comparing the predicted to actual PMIs with the testing dataset was used to assess the quality of the prediction (the fit (i.e., R^2^) and slope.

**Gene annotation**. The genes were annotated by performing BLASTN searches using the NCBI databases. Genes that had a bit score of greater than or equal to 100 were annotated.

**Experimental design**. Three different datasets were used in this study: the whole zebrafish transcriptome, the mouse brain transcriptome, and the mouse liver transcriptome. The datasets were split into training and testing data. The training data was used to build the regression equations and the testing data was used to validate the equations. Three different experimental designs were tested.

1. Simple linear regressions using individual genes. We examined if simple linear regressions (*PMI*_predict_=*m*\* transcript abundance + *b*) of individual gene transcripts could be used to predict PMIs. The values of *m* and *b* were determined using the training dataset. The performance of the regression was assessed using the R^2^ of the predicted versus actual PMIs with both training and testing datasets.
2. Over-defined linear regressions using top performing genes from Experimental Design 1. An over-defined linear regression is used when the data consisted of more rows (postmortem times) than columns (gene transcripts). The top three individual gene transcripts in Experimental Design 1 were combined and trained to predict PMIs using an over-defined linear regression model. The performance of the model was assessed using the R of the predicted versus actual PMIs of both training and testing datasets.
3. Perfectly defined linear regressions using randomly selected gene transcript sets. A perfectly-defined linear regression is used for data consisting of equal number of rows (postmortem times) and columns (gene transcripts). A random number generator was used to select sets of genes from the datasets in order to find the top PMI predictors. The analysis yields a set of coefficients (i.e., *m’s*), one coefficient for each gene transcript in a set. The coefficients were used to predict the PMIs of a gene set. The R^2^ and slope of the predicted versus actual PMIs were determined using the training and testing data. The procedure of selecting the gene transcript sets from the training set, determining the coefficients, and testing the coefficients was repeated at least 50,000 or more times and the gene transcript sets generating the best fit (R^2^) and slopes were identified (Fig 1).

**Fig 1.**
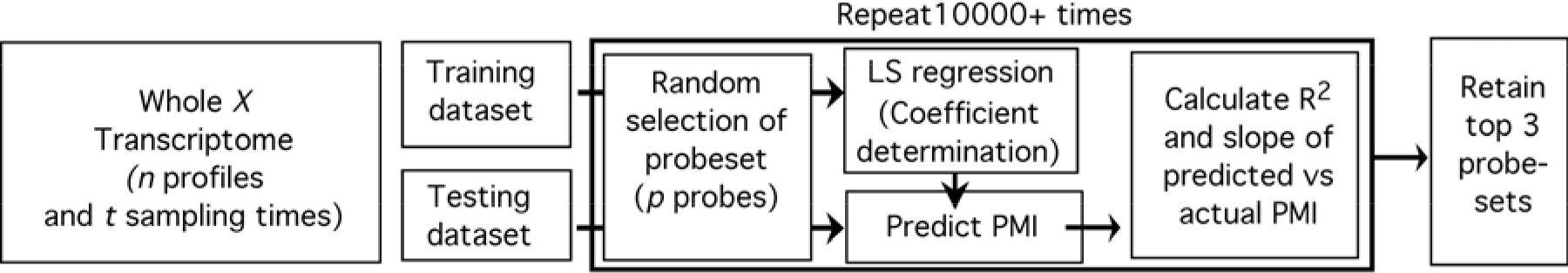
Cartoon of experimental design for three different datasets. Bold box was repeated 10,000+ times. The top 3 probe datasets were determined by the R^2^ between predicted versus actual PMI and the slope closest to one using the test dataset. If X=’zebrafish’ then *n*=548, *t*=11, p=11; if X=’mouse brain’ then **n**=478, *t*=7, p=7; if X=’mouse liver’ then *n*=36, *t*=7, *p*=7.

## Results

The 36,811 probes of the zebrafish and 37,368 probes of the mouse were calibrated. Of these, the transcriptional profiles of 548 zebrafish genes and 515 mouse genes were found to be significantly upregulated. Of the 515 upregulated genes, 36 were from the liver and 478 genes were from the brain. It is important to note that each datum point in a zebrafish transcriptional profile represents the mRNA obtained from two zebrafish and each datum point in a mouse profile represents the mRNA obtained from one mouse. In other words, each datum point represents a true biological replicate. Duplicate samples were collected for each postmortem time for the zebrafish profiles, and triplicate samples were collected for the mouse (with exception of the 48 h postmortem sample which was duplicated) at each postmortem time.

### Predicting PMI with 1 or 3 gene transcripts

The ability of individual gene transcripts to accurately predict PMIs was assessed using the simple linear regression: 
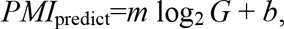
with *m* as the slope (i.e., the coefficient), *G*is the individual gene transcript abundance, and *b* is the intercept.

For the zebrafish, one of the duplicates (at each postmortem time) was used to determine the linear regression equation (i.e., *m* and b) and the other one was used to test the regression equation. For the mouse, one of the triplicates at each postmortem time was used to determine the linear equation and the remaining data (2 data points) were used to test the regression equation. The three gene transcripts of the zebrafish, mouse brain, and mouse liver with the highest fits (R^2^) between predicted and actual PMIs are shown in Table 1.

**Table 1.** Top three fits (R^2^) of predicted and actual PMIs by organism/organ based on the training and testing datasets of individual probes (probe names were designated by Agilent) targeting specific transcripts. Corresponding gene names and functions are shown. Whole, RNA was extracted from whole organisms; Brain, RNA extracted from mouse brains; Liver, RNA extracted from mouse livers.

| Organism/<br>Organ | Oligonucleotide<br>Probe Name | $R^2$ | Number of data points | Gene Name and Function |
| --- | --- | --- | --- | --- |
| Zebrafish |  |  |  |  |
| Whole | A_15_P121158 | 0.94 | 11 duplicates | Non-coding |
|  | A_15_P295031 | 0.84 | 11 duplicates | Non-coding |
|  | A_15_P407295 | 0.82 | 11 duplicates | Non-coding |
| Mouse |  |  |  |  |
| Brain | A_66_P130916 | 0.67 | 6 triplicates and 1 duplicate | Histocompatibility 2, O region beta locus |
|  | A_55_P2127959 | 0.61 | 6 triplicates and 1 duplicate | Zinc finger protein 36, C3H type-like 3 |
|  | A_55_P2216536 | 0.60 | 6 triplicates and 1 duplicate | E3 ubiquitin-protein ligase |
| Liver | A_55_P2006861 | 0.94 | 6 triplicates and 1 duplicate | Triple functional domain protein |
|  | A_30_P01018537 | 0.91 | 6 triplicates and 1 duplicate | Prokineticin-2 isoform 1 precursor |
|  | A_51_P318381 | 0.90 | 6 triplicates and 1 duplicate | Placenta growth factor isoform 1 precursor |

For the zebrafish, the gene transcript targeted by probe A_15_P121158 yielded a fit (combined training and testing data) of R^2^ =0.94, while the other gene transcripts yielded moderate fits (R^2^<0.90). The top predictors of PMIs for the mouse brain samples yielded weak R^2^-values (0.61 to 0.67), and the top predictors for the mouse liver samples yielded reasonable R^2^-values (0.90 to 0.94) (Table 1) suggesting that the liver was more suitable for predicting PMI than the brain.

In addition to assessing the PMI prediction of individual gene transcripts, we investigated if a combination of the top gene transcripts would improve upon PMI predictions. Using an over-defined linear regression: 
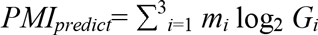
and one of the duplicate/triplicate samples from each postmortem time as the training data, we determined the coefficient for each gene transcript and tested the regression equation using the remaining test data. For the zebrafish, the derived coefficients for genes targeted by probes A_15_P295031, A_15_P121158, and A_15_P407295 were −162.97, 22.44, and 35.10, respectively. Using the gene transcript abundances for these probes at 48 h postmortem (−0.33 a.u., −0.89 a.u., and −0.25 a.u., respectively) and the equation above, the predicted PMI is ~25.3 h. For the mouse brain gene transcripts targeted by probes A_66_P130916, A_55_P2127959, and A_55_P2216536, the derived coefficients were 3.70, −3.57, and 45.25, respectively. Using the gene transcript abundances for these probes at 48 h postmortem (1.21 a.u., 0.36 a.u., and 0.80 a.u., respectively) and the equation above, the predicted PMI is ~39.2 h. For the mouse liver gene transcripts, the derived coefficients targeted by probes A_51_P318381, A_30_P01018537, and A_55_P2006861 were −3.75, 36.21, and −13.93, respectively. Using the gene transcript abundances for these probes at 48 h postmortem (1.04 a.u., 1.65 a. u., and 0.48 a.u., respectively) and the equation above, the predicted PMI is ~49.3 h. The fits (R^2^) of the predicted versus actual PMIs for the zebrafish, the mouse brain and mouse liver were 0.74, 0.64, and 0.86, respectively.

While some of the individual gene transcript abundances yielded reasonable PMI predictions using simple linear equations (Table 1), combining the individual gene transcript abundances and using an over-defined linear regression did not significantly improve upon PMI predictions based on individual genes.

These experiments showed that neither simple linear regression equations derived from the individual gene transcripts, nor over-defined linear regressions derived from the top three individual gene transcripts satisfactorily predicted PMIs.

### Predicting PMI with many genes

To predict PMIs using perfectly defined linear regressions, the number of gene transcripts used for the regression has to equal the number of postmortem sampling times. The zebrafish was sampled 11 times and the mouse was sampled 7 times, therefore 11 and 7 genes could be used for the regressions, respectively. The regression equation for the zebrafish was: 
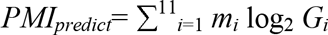

The regression equation for the mouse was: 
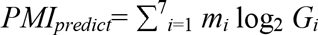

The procedure to find gene transcript sets that provide the best PMI predictions included: assigning randomly-selected genes to gene transcript sets, determining the coefficients of the gene transcripts in the set using a defined least squared linear regression, and validating the regression model with gene transcript sets in the test data. We rationalized that if this process was repeated thousands to millions of times, groups of gene transcripts could be identified that accurately predict PMIs with high R^2^=<0.95 and slopes of 0.95 to 1.05.

The number of upregulated genes in the zebrafish, mouse brain, and mouse liver transcriptome datasets is relevant to determining the optimal gene transcript set for PMI predictions because of the magnitude of possible combinations to be explored. For example, there are 2.85 × 10^22^ combinations of 11 gene transcripts for the zebrafish dataset (*n*=548), 1.08 × 10^15^ combinations of 7 gene transcripts for the mouse brain dataset (*n*=478) and 8.35 × 10^6^ possible combinations of 7 gene transcripts for the mouse liver dataset (n=36). Therefore, for some transcriptome datasets (i.e., zebrafish and mouse brain), the determination of the best gene transcript set to accurately predict PMI was constrained by the number of possible combinations explored in reasonable computer time.

### Validation of PMI prediction

After training 50,000 random selections, about 95% (*n*=47,582) of the selected gene transcript sets yielded R^2^ and slopes of 1 with the training datasets. The remaining selections (*n*=2,418) did not yield R^2^ and/or slopes of 1 because the equations could not be resolved, or else the fits and/or slopes were <1. The R^2^ and slopes of the predicted versus actual PMIs using the testing data were used to identify the top gene sets.

The top three gene transcript sets with the highest R^2^ and slopes closest to one are shown in Fig 2. The gene transcript set used in Panel A had smaller confidence intervals than those found in Panels B and C. At the 99% confidence level, the predicted PMIs for the gene set in Panel A ranged from 7 to 11 h for the actual PMI of 9 h, from 8 to 16 h for the actual PMI of 12 h, from 21 to 27 h for the actual PMI of 24 h, from 46 to 50 h for the actual PMI of 48 h, from 96 h for the actual PMI of 96 h. These results suggest that PMIs could be accurately predicted using zebrafish gene sets and the derived coefficients.

**Fig 2.**
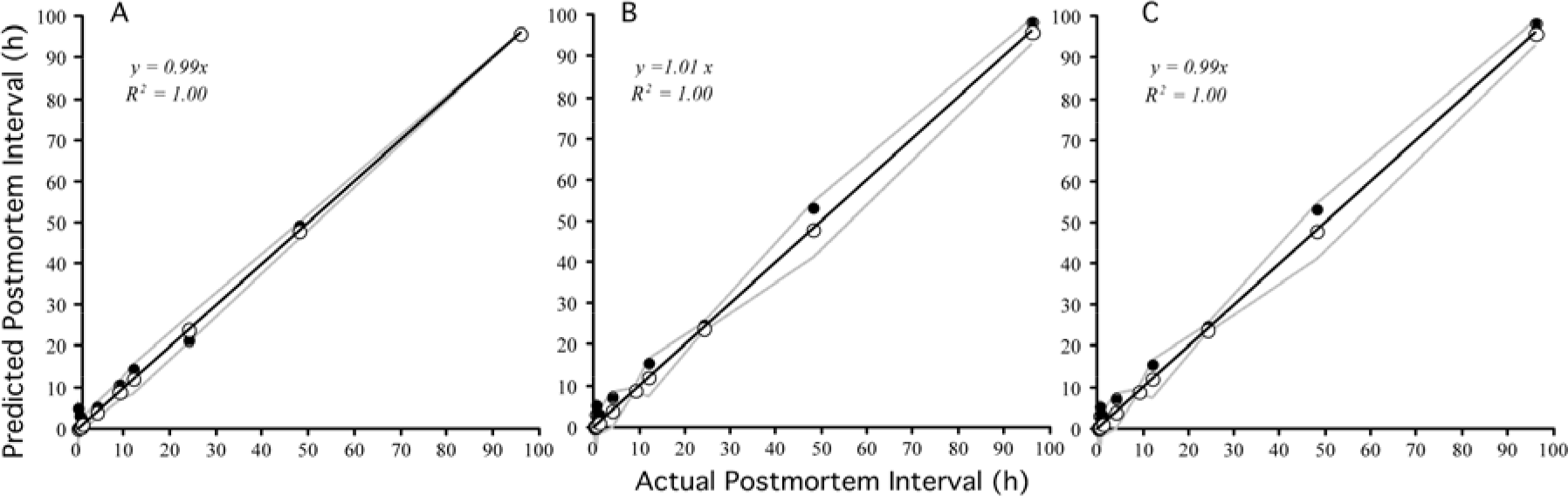
Predicted versus actual PMIs determined for the zebrafish by three equations representing different gene transcript sets. R^2^ and slopes are based on both training and testing datasets. Gray line represents 99% confidence limits of the linear regression. Open circles, training data; closed circles, testing data. See Table 1 for information on the genes and annotations.

The gene transcription profiles of the 11 genes used in Fig 2, Panel A are shown in Fig 3. Note that the gene transcript abundances of the duplicate samples used for training and testing are similar at all sampling times. These results show the high precision of the Gene Meter approach since each datum point represents different zebrafish. Note that each gene has a different postmortem transcriptional profile.

**Fig 3.**
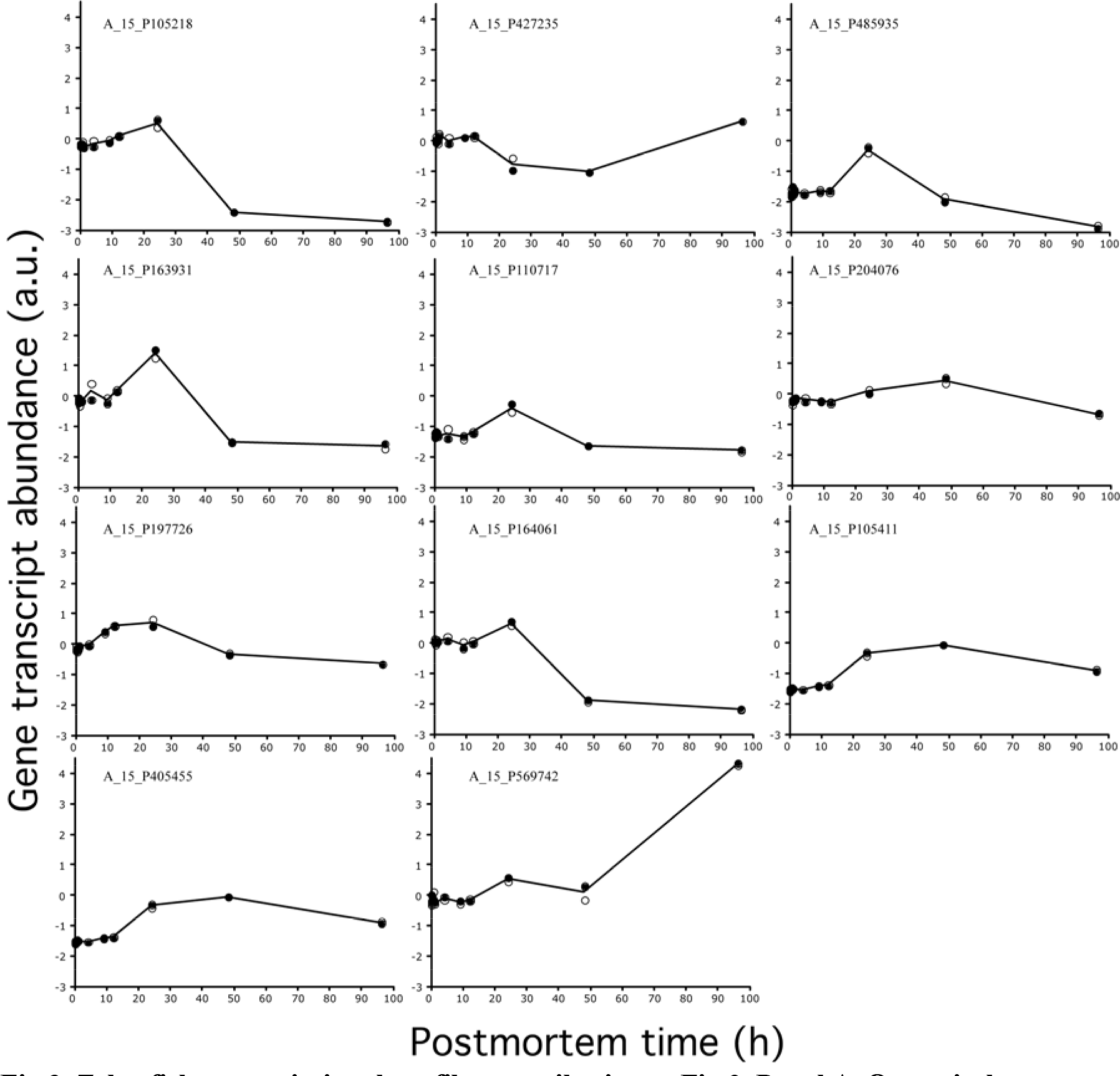
Zebrafish transcriptional profiles contributing to Fig 2, Panel A. Open circles, training data; closed circles, testing data; black line, average. See Table 2 for information on the equations and probe annotations.

Table 2 shows the probe labels for the gene transcript sets used in Fig 2 Panels A to C and their corresponding coefficients derived from the training dataset. Note that only some genes could be annotated using NCBI. We assumed that genes not annotated represent non-coding mRNA. The PMIs in Panel A to C could be predicted by adding the products of the log_2_ abundance of each gene to its corresponding coefficient. For example, the equation for Table 2 Panel A is: 
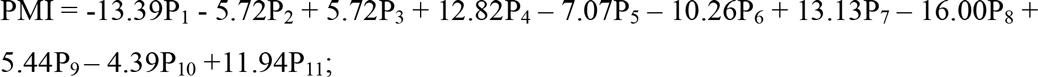
where P_i_ are the gene abundances represented by the probes A_15_P105218 (0.39 a.u.), A_15_P427235 (−0.54 a.u.), A_15_P485935 (−0.39 a.u.), A_15_P163931 (1.27 a.u.), A_15_P110717 (−0.53 a.u.), A_15_P204076 (0.16 a.u.), A_15_P197726 (0.82 a.u.), A_15_P164061 (0.58 a.u.), A_15_P105411 (−0.40 a.u.), A_15_P405455 (−1.13 a.u.), A_15_P569742 (0.46 a.u.). In this example, the predicted PMI is ~24 h. Based on Figure 2 panel A, the 99% confidence interval is between 20.9 and 27.1 h.

**Table 2.**
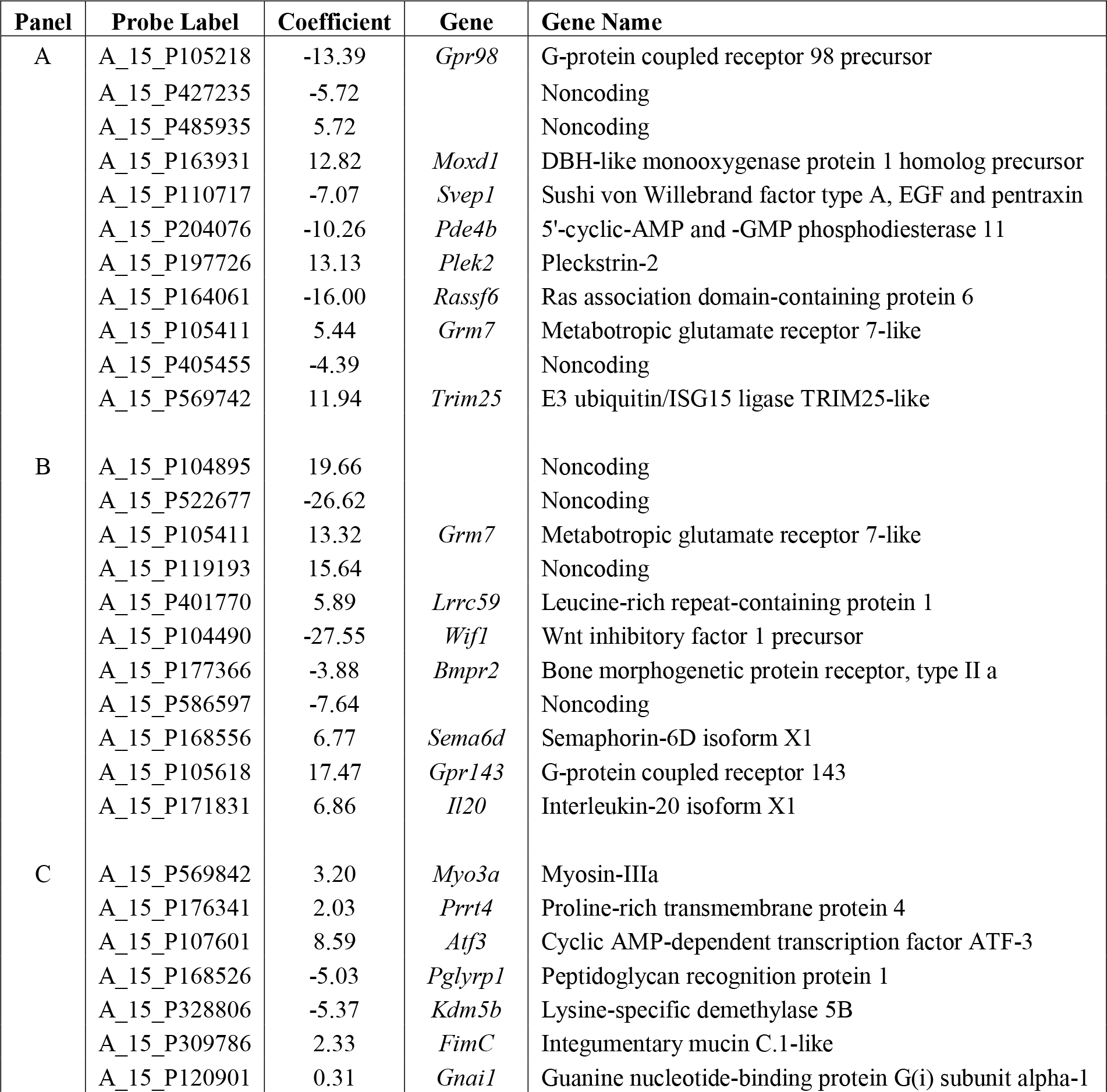
Zebrafish genes used to predict PMI by Panel. The gene annotations of the probes were determined using NCBI with a 100 bit minimum.

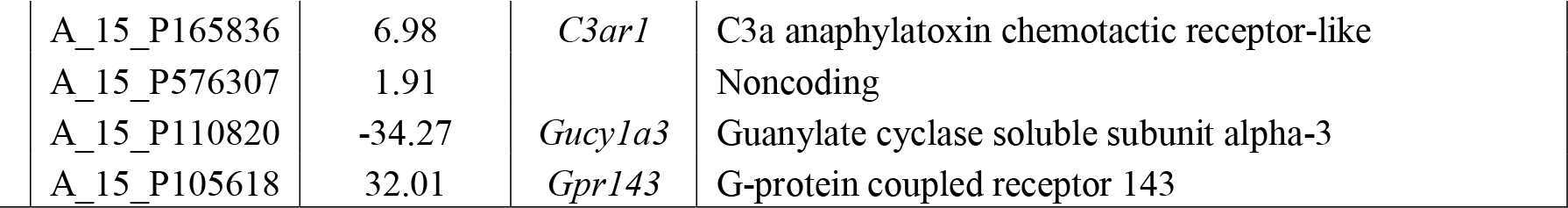

### Mouse

After training 50,000 random selections (each selection consisted of 7 genes), about 96% (*n*=47,847) of the selected gene sets yielded R^2^ and slopes of 1. The remaining selections were not used for validation because the equations (*n*=2,153 selections) could not be resolved, or they had fits and/or slopes that were <1 (*n*=25 selections). The R^2^ and slopes of predicted versus actual PMIs determined using the testing dataset identified the top performing gene sets.

The top selected gene transcript sets for the mouse liver and brain are shown in Fig 4. As indicated by the R^2^, slopes, and size of the 99% confidence intervals, gene transcript sets from the liver were better at predicting PMIs than those from the brain. The mouse genes used in the gene transcript sets, their coefficients, and annotations are shown in Table 3 and the transcriptional gene profiles for the mouse liver samples are shown in Fig 5. Note that the high similarity in the gene transcript abundance between the data used for training and testing of the selected genes. In most cases (but not all), the duplicate samples (represented by dots) are located on top of one another.

**Fig 4.**
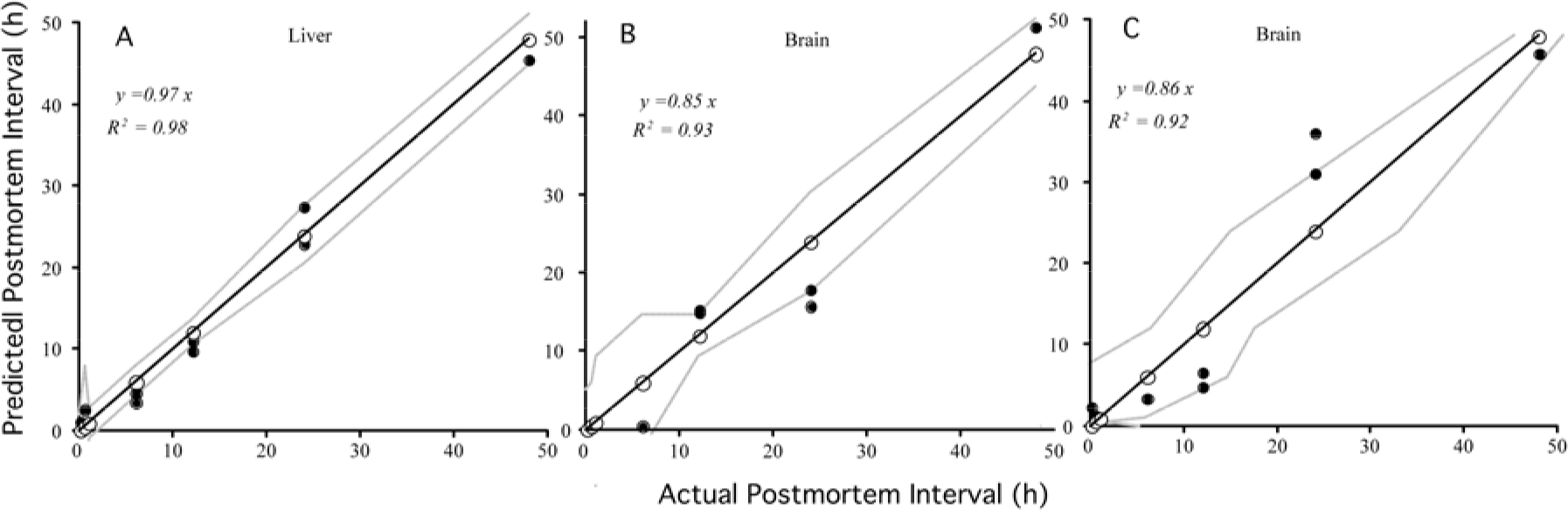
Predicted versus actual PMI determined for the mouse for three different equations as represented by the panels. R^2^ and slopes are based on both training and testing datasets. Gray line represents 99% confidence limits of the linear regression. Open circles, training data; closed circles, testing data. See Table 3 for information on the equations and probes.

The poor predictability of the brain gene transcript sets (i.e., R^2^<0.95) could be attributed to the low number of repeated selections of gene transcript sets and the variability in gene abundances between the training and testing datasets. We repeated the analysis of the brain samples an additional 1,000,000 times, which resulted in some improvement. The best fit and slope for 50,000 gene transcript set selections was R^2^ =0.83 and *m*=0.77 (not shown). The best fit and slope for 1,000,000 selections was R^2^=0.93 and *m*=0.85 (Fig 4, Panel B) with the second best being R^2^=0.92 and *m*=0.86 (Fig 4, Panel C). Hence, the number of combination of gene transcript sets examined is important for selecting the best ones. It is important to emphasize that the computation time for running 1,000,000 selections was approximately 1 week using a Mac OS X 10.8.6.

The PMIs in Panel A to C could be predicted by adding the products of the log_2_ abundance of each gene to its corresponding coefficient. The predicted PMIs for mouse is calculated same way as for zebrafish (shown above).

**Table 3.**
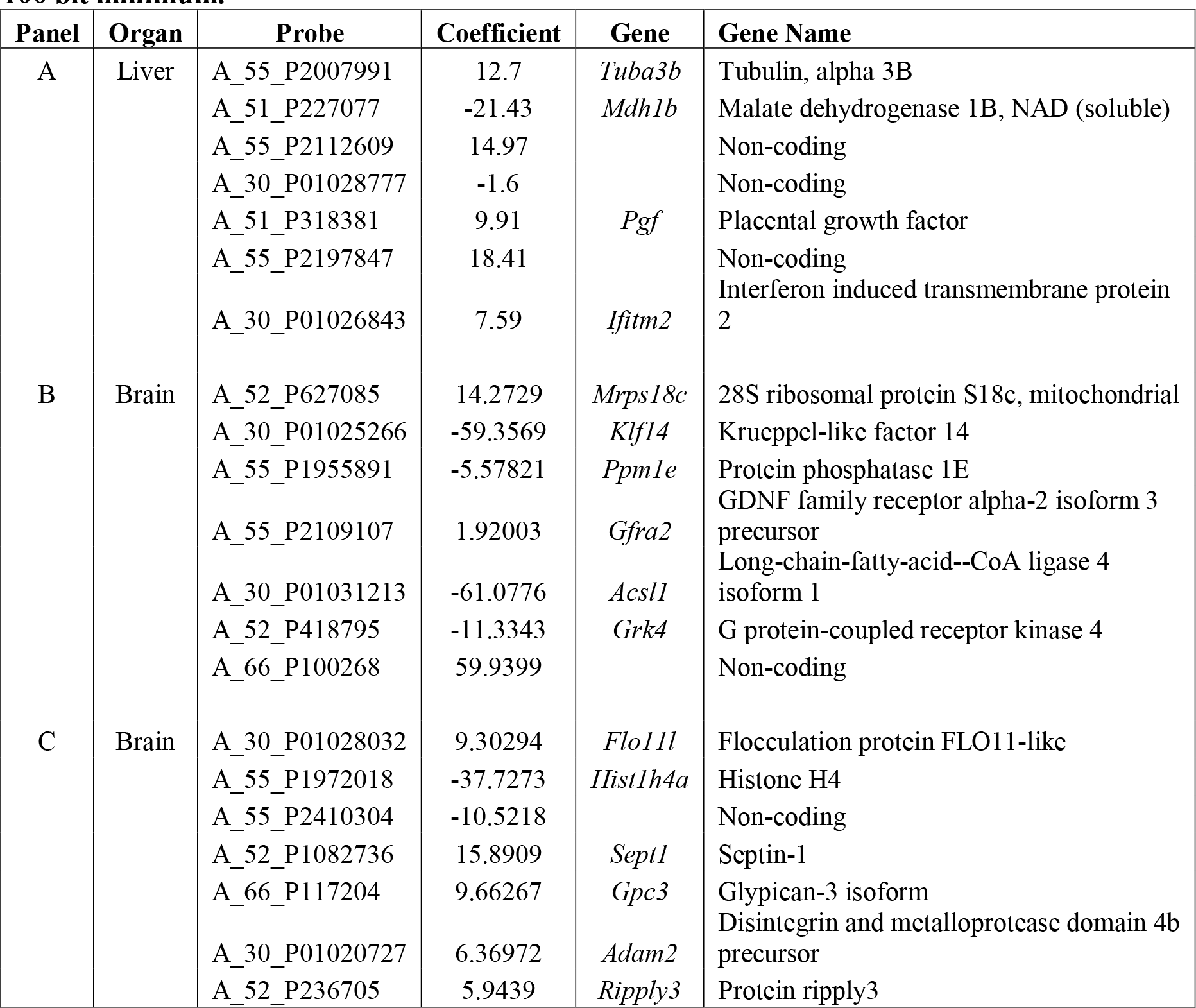
Mouse probes used to target gene transcripts and their coefficients used to predict PMIs by Panel. The annotations of probes were determined by using NCBI database with 100 bit minimum.

We compared the variability in gene transcript abundances between training and testing data sets for the mouse liver and mouse brain. Transcriptional gene profiles of the gene sets used in Fig 4 Panels A and B are shown in Figs 5 and 6, respectively. While most of the mouse liver gene transcript abundances are similar for the training and testing data sets in Fig 5, many of the mouse brain gene transcript abundances are not similar in Fig A two-tailed T-test of the standard deviations of the gene transcript abundances in the training and testing datasets for the liver and brain samples (Fig 5 versus Fig 6) by postmortem time were significantly different (P< 0.006), with higher standard deviations in the brain samples than the liver. This finding indicates that variability in the gene transcript abundances affects PMI predictability.

**Fig 5.**
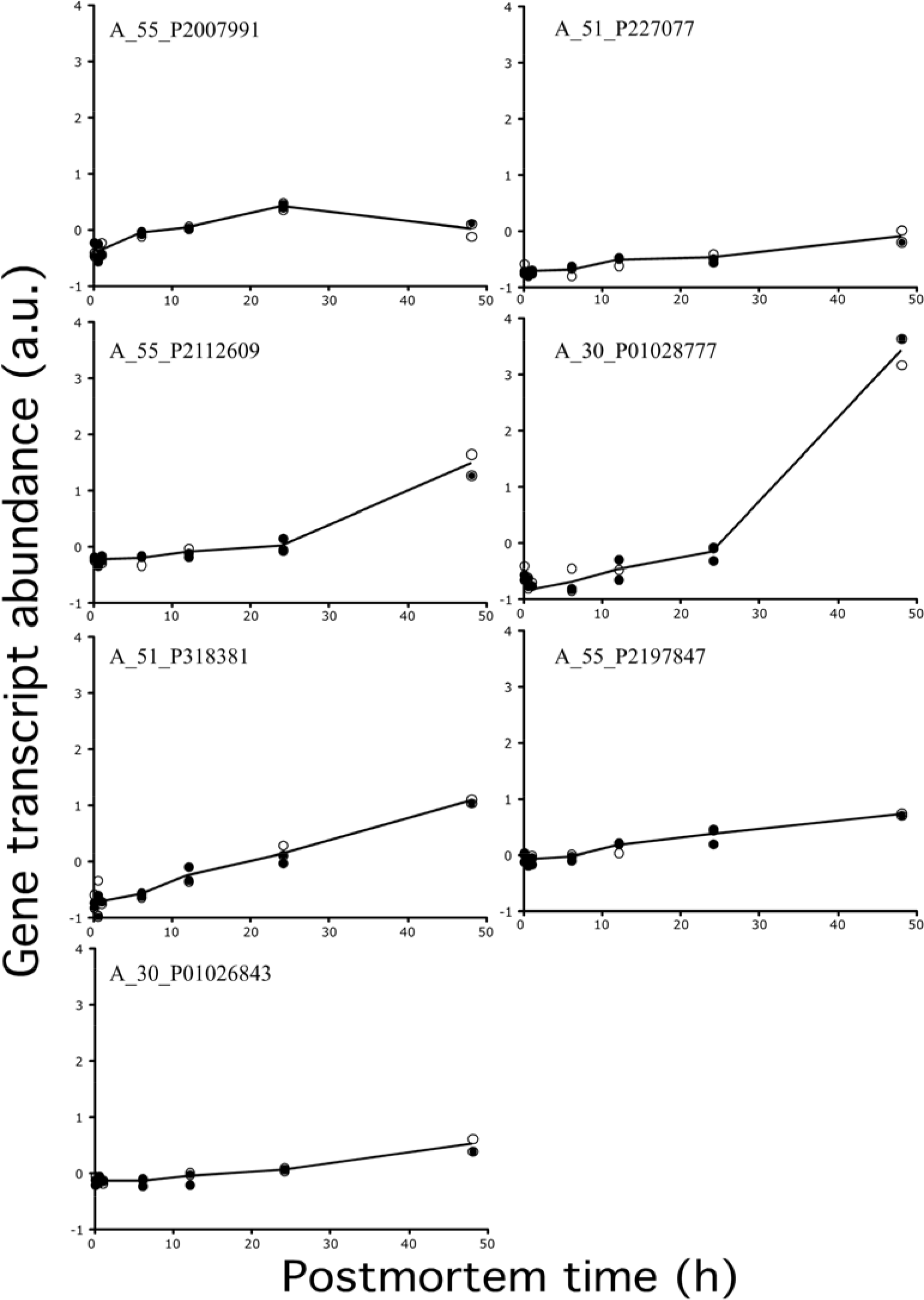
Mouse liver transcriptional profiles contributing to Fig 4, Panel A. Open circles, training data; closed circles, testing data; black line, average. See Table 3 for information on the equations and probe annotations.

**Fig 6.**
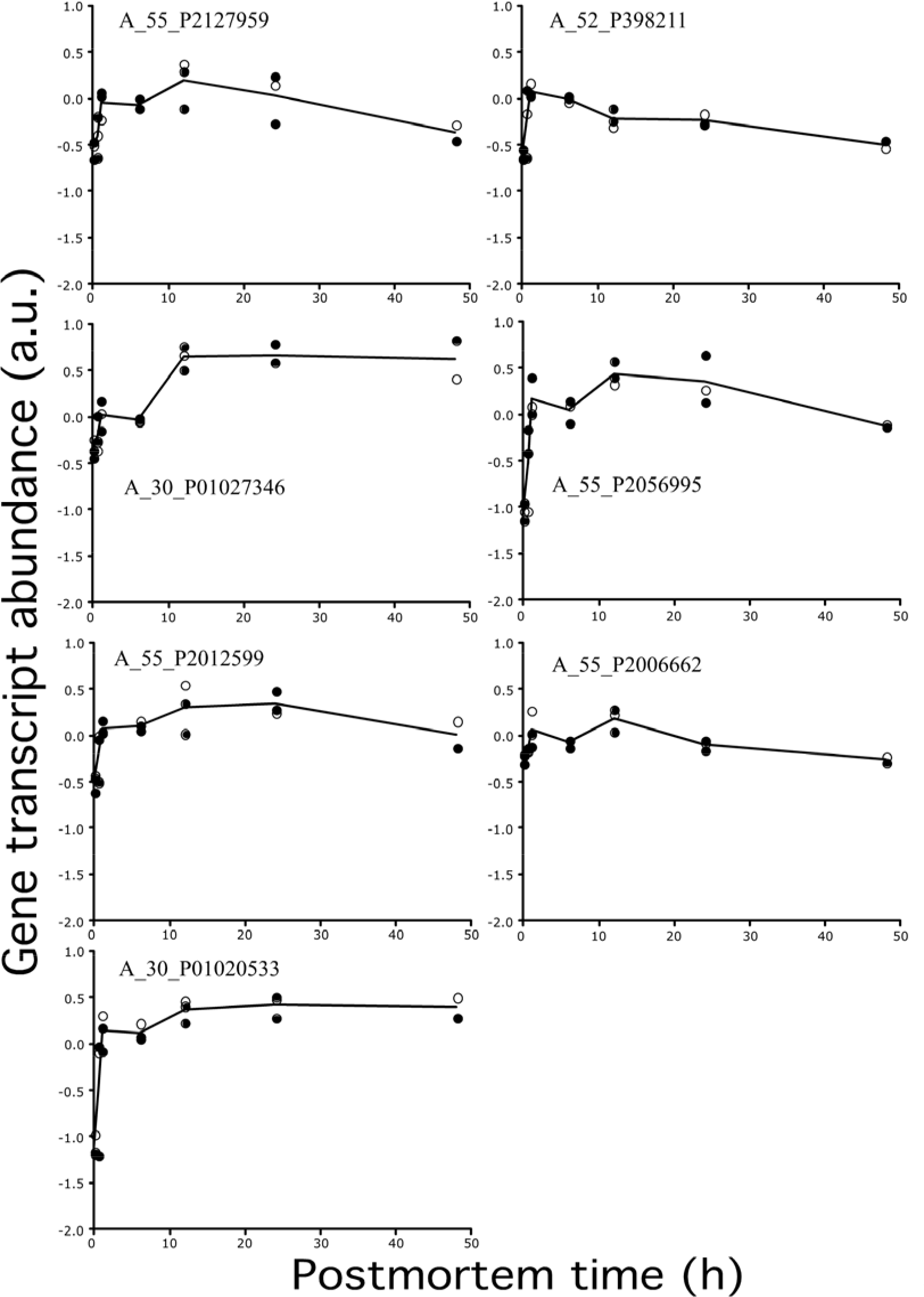
Mouse brain transcriptional profiles contributing to Fig 4, Panel B. Open circles, training data; closed circles, testing data; black line, average. See Table 3 for information on the equations and probe annotations.

To further test this phenomenon, a small amount of random noise was added to the abundances of mouse liver gene transcripts (Fig 5), which originally had very low standard deviations by postmortem time. When the introduction of noise approached 10%, the fit and slopes were drastically altered, indicating that similarity in the gene transcript abundances between the training and test data sets can directly affect the fit and slopes of predicted versus actual PMIs.

### Randomization challenge

Experiments using perfectly-defined linear regressions revealed that 95% of the training data for the zebrafish and mouse yielded fits (R^2^) and slopes of 1 for predicted versus actual PMIs. To demonstrate that the ‘perfect’ fits and slopes are functions of the linear regressions, we randomized gene transcript abundances at every postmortem time for all genes in the zebrafish dataset. This randomization maintained the variance in the dataset at each postmortem time so that the variance of the first postmortem time in the randomized dataset was the same as the variance of the gene transcript abundances at the first postmortem time in the original dataset and so on for the gene transcript abundances at all postmortem times.

As anticipated, training of the randomized zebrafish dataset using perfectly-defined linear regressions (11 genes by 11 postmortem times × 50,000 repeated gene transcript selections) yielded fits (R^2^) and slopes of 1 for predicted versus actual PMIs for 95% of the regressions. When we tested the 50,000 regressions using a test dataset, not one of the gene transcript sets approached a fit (R^2^) and slope of 1. In fact, most yielded slopes of zero and R^2^ <0.80. The significance of this experiment is twofold: (i) it confirms that ‘perfect’ fits and slopes using the training datasets are a function of ‘perfectly-defined’ linear regressions, and (ii) it confirms the need to validate the regression equations using testing datasets.

## Discussion

In addition to the different stages of body decomposition (i.e., rigor mortis, livor mortis, algor mortis and putrefaction) [16,17,18], there are many biochemical, biological, chemical, and physical ways to determine PMI. Biochemical indicators and corresponding sample sites include: pH and spectrophotometer readings of blood and serum [19], cardiac troponin-I and cadaveric blood in the heart [19,20], lactate and malate dehydrogenase in the liver [21], melatonin in the brain, sera, and urine [22], DNA degradation in many tissues and organs [23,24,25,26], endothelial growth factors in the brain, heart, liver, and kidneys [27], insulin and glucagonin in pancreatic beta cells [28,29], cells in the cerebrospinal fluid [30], apoptotic cells in skin bruises [31] and histology of labial muscosa [32]. Biological indicators and sample sites include: ciliary motility in the nose [33], sweat gland morphology in the arm pit [34], muscle contraction [35] and pyrosequencing of the buccal cavity, rectum and GI tract samples [36], entomological [37,38,39] and botanical processes occurring in and around the body [40,41]. Chemical indicators and sample sites include: electrolytes in human vitreous humour [42], biomarkers (e.g. amino acids, neurotransmitters) in body organs and muscles [43], hypoxanthine in the vitreous humour or cerebrospinal fluid [44,45,46] and potassium in the vitreous humour [47,48,49]. Physical indicators and sample sites include: microwave probe to the skin [50], infrared tympanic thermography and temperature of the ear [51,52], and temperature of the eye and body core [53,54,55]. Several formulae have also been developed to estimate PMI that are based on multiple environmental and physicochemical conditions [e.g., 56]. Despite these many approaches, accurate PMI prediction remains an enigma [43]. The motivation for this study was to test experimental designs that could accurately predict PMI using upregulated gene expression data in order to provide “proof of principle”.

The abundance of a gene transcript is determined by its rate of synthesis and its rate of degradation [57]. In this study, the synthesis of mRNA had to far exceed its degradation to be a significantly upregulated gene (at some postmortem time) in our study. As demonstrated in the previous study [13] and shown in this study (Figs 3, 5, and 6), the timings of the upregulation differed between genes. Some gene transcripts, such as the one targeted by probe A_15_P105218, were upregulated right after organismal death and reached maximize abundance at 24 h postmortem while, others, such as the one targeted by probe A_15_P569742, increased substantially at 48 h postmortem (Fig 3). It is presumed that differences in the transcript profiles affect the value of the coefficients in the linear equations because it is not possible to generate coefficients if the gene transcript abundances changed in the same way. That is, a numerical solution could not be mathematically resolved.

It should be noted that the upregulation of postmortem genes is optimal for PMI prediction because only about 1% of the total genes of an organism were upregulated in organismal death - which is rare indeed. In contrast, a focus on downregulated genes would not be practical because one does not know if downregulated genes are due to repression, degradation of the total RNA, or exhausted resources such as those needed for the transcript machinery function (e.g., dNTPs and RNA polymerase).

Given that gene transcripts from the liver were better at PMI predictions than those from the brain suggests that mRNA transcripts from some organs are better than others. It is conceivable that upregulated postmortem genes could be found in the heart, kidney, spleen or muscle, which needs further exploration.

It is important to recognize that this study would probably not be possible using conventional microarray approaches because normalizations could yield up to 20 to 30% differences in the up-or down-regulation depending on the procedure selected [59–62]. The Gene Meter approach does not require the data to be normalized since the microarray probes are calibrated. Moreover, in the processing of samples, the same amount of labeled mRNA was loaded onto the DNA microarray for each sample (1.65 μg), which eliminates the need to divide the microarray output data by a denominator in order to compare samples.

We recognize that our experimental design did not consider factors such as temperature, which have been considered in other models [e.g., 4]. To do so would go beyond the stated objectives of providing a “proof of principle” for the optimal experimental design (i.e., perfectly-defined linear regressions based on multiple gene transcripts) using a high throughput approach. Nonetheless, future studies could make our experimental design more universal by integrating temperature and other factors into the regression models.

In addition to providing “proof of principle” of a new forensic tool for determining PMI, the approach could be used as a tool for prospective studies aimed at improving organ quality of human transplants.

## Conclusion

We examined if significantly upregulated genes could be used to predict PMIs in two model organisms using linear regression analyses. While PMIs could be accurately predicted using selected zebrafish and mouse liver gene transcripts, predictions were poor using selected mouse brain gene transcripts, presumably due to high variability of the biological replicates. The experimental design of selecting groups of gene transcripts, extracting the coefficients with linear regression analyses, and testing the regression equations with testing data, yielded highly accurate PMI predictions. This study warrants the implementation of our experimental design towards the development of an accurate PMI prediction tool for cadavers and possibly a new tool for prospective studies aimed at improving organ quality of human transplants.

## Authors’ contributions

PAN and AEP designed the study. MCH and PAN conducted the statistical analyses and wrote the manuscript. All authors read and approved the final manuscript.

## Supplementary Information

### Primer on Matrix Algebra

Our PMI prediction approach using a perfectly defined system relies on matrix linear algebra. The following is an explanation of matrix linear algebra and how it was used in our study to predict PMI with gene expression profiles. A matrix is defined as a rectangular array of related values. These values, which are called elements, usually are scalars. Scalars are numbers that represent physical quantities. Elements in a horizontal line are called rows and elements in a vertical line are columns and the number of rows and columns describe a matrix. A matrix, **S**, with *y* number of rows and z number of columns is denoted as a *y* × z matrix. This matrix can also be notated with subscripts and appears as **S**_y,z_. Mathematical operations can be performed using matrices, including multiplication. Matrices can be multiplied by one another if one matrix has as many columns as the other matrix has rows. We used linear equations to obtain a matrix product. The number of gene transcripts used in a selected gene transcript set was limited by the number of postmortem sample times.

### Example

If we arrange individual genes with their transcript abundances at specific postmortem times in columns then, essentially, we have a matrix **A** where the columns are the transcriptional profiles for individual genes. Furthermore, we can construct another matrix, **B**, which defines the data in a different way. In this matrix the values are the actual postmortem sampling times that are ordered in the same way as the abundances of individual gene transcripts. Finally, we can define another matrix, consisting of one column of coefficient values. These are the weighing factors that we will determine using the rules of linear matrix algebra. To deconvolute the transcriptional profiles from the mixtures and solve for the weighing factors, we set up the matrices like so: **A** × C = **B**. When this is done we are left with several equations. To solve for *x* and *y* we need to follow the rules of linear matrix algebra. First, we must transpose **A**, which becomes A^T^, then multiply both sides of the equation **A** × **B** = **C** with **A**^T^ Next, we are required to invert the matrix product of **A** × **A**, thus **A** becomes **A**^−1^ and also multiply both sides of our original equation **A** × **C** = **B**. Our modified equation, which looks like this (**A**^T^ * A)^−1^**A**^T^*A\***C** = (**A**^T^*A)^−1^ * **A**^T^ * **B**, is now ready to be solved for *x* and *y* in matrix **C**. Solving for **C**, we get C = (**A**^T^ * **A**)^−1^ * **A**^T^ * **B**. Now the values in the matrix **C** are *x* and *y*. Next, we plug our values for *x* and *y* into our matrix equations to obtain predicted (calculated) PMI values. To ascertain whether the predicted PMIs from our group of gene transcripts is accurate, we plotted the actual and predicted PMIs and determined the slope and fit of the regression line. The R^2^ and the slope of the line was observed to determine how well a group of gene transcripts predicted PMI. The R is the measure of how much variability is accounted for by the model. For example, if the R^2^ is 0.95, then the model accounts for 95% of the variability. The other 5% is due to undetermined phenomena.

